# Fungicide and warming interact to reduce soil ecosystem functioning

**DOI:** 10.64898/2026.01.16.699873

**Authors:** Matthew Kelbrick, Jake McEntagart, Edward Cairns, Steve Paterson, Sharon E. Zytynska, James P. J. Hall, Siobhan O’Brien

## Abstract

Soil microbial communities are essential to the functioning of their ecosystems, providing vital services such as plant growth promotion and bioremediation. However, intensively managed environments such as agricultural soils are increasingly challenged with multiple harsh environmental stressors from the compounding effects of climate change and conventional agricultural practices. Understanding how soil communities will respond to environmental stress along multiple axes is key for predicting how environmental change will impact terrestrial soil ecosystems. We combine experimental evolution with community phenotyping, 16s rRNA metabarcoding, fungicide resistance assays and plant experiments, to show that fungicide and warming temperatures together drive a synergistic reduction in soil community respiration rates, metabolic capacity and plant growth promotion (*Hordeum vulgare*), which would have been missed by looking at stressor effects independently. Dual stressors caused widespread loss of metabolic activity, particularly for carbohydrates and carboxylic acids, highlighting impairment of community function. Fungicide-evolved soil isolates had a 16-fold increase in fungicide resistance, but only in the absence of warming – suggesting that the dual stressor treatment constrained fungicide adaptation. Together, our findings suggest that the number of stressors (rather than the nuances of individual stressors) may be key for predicting soil microbial community responses to environmental change across multiple axes.

## 1. Introduction

Soils harbour diverse and complex microbial communities that perform essential ecological functions, including nutrient cycling, carbon storage and crop protection [1]. These activities are increasingly threatened by anthropogenic activities such as pesticide application, chemical pollution, and land-use change, which can alter soil community composition, reduce microbial diversity, and impair ecosystem functioning [2–7]. Although the impacts of individual stressors on soil communities are often studied in isolation, soil environments are frequently subjected to multiple, simultaneous disturbances [4, 8]. For example, climate change-induced heatwaves may co-occur with agricultural practices known to cause fungicide and antibiotic runoff. Mounting evidence suggests that the combined effects of multiple stressors can yield non-additive, often unpredictable outcomes that differ from single-stressor scenarios [4, 8–11]. Despite this understanding, the vast majority of studies addressing the impact of global change on microbial communities tend to examine global change factors individually [8].

Despite growing awareness of anthropogenic pressures on terrestrial ecosystems, the interactive effects of multiple stressors on soil microbial communities, and consequences for microbial ecosystem functioning, remain poorly understood. Addressing this knowledge gap could provide information on which stressors to prioritise for maintaining ecosystem stability under climate change [12]. When multiple stressors act on a community, their combined effects can be synergistic (combined effects are more than expected), antagonistic (combine effects are less than expected) or additive (combined effects are equal to the sum of individual effects; i.e. the null model) [13]. For instance, species that tolerate one stressor may differ from those that can withstand another, and few may be capable of tolerating both. Consequently, when stressors are combined, the community may lack taxa able to persist under the joint conditions [14], so that even additive effects on species composition can have synergistic effects on functioning. Over longer timescales, evolutionary trade-offs and antagonistic pleiotropy could also underlie synergistic stressor effects [13], while cross-resistance, co-resistance, or co-regulation could underlie antagonistic effects of multiple stressors [15, 16]. Multiple stressors can also reduce the evolutionary potential of populations by lowering population sizes and inducing genetic bottlenecks, which limit genetic variation and the supply of beneficial mutations (Hiltunen et al., 2015).

Pesticides are widely used in agriculture to maintain sufficient food production amid global population growth. In 2021, 355,175 tonnes of pesticides were sold in the European Union [17]. However, only a fraction reaches target pests; estimates suggest that 30–50% of applied pesticide is lost by deposition on the ground, or volatized [18, 19]. Pesticides may exert unintended effects on non-target organisms by targeting cell membrane components, protein synthesis, signal transduction, respiration, cell mitosis, and nucleic acid synthesis [20, 21]. Pesticides narrow microbial metabolic breadth, select for pesticide degraders and resistant strains, and accelerate nutrient loss from soils, especially when multiple classes of pesticides are applied simultaneously [5]. Studies investigating the impact of pesticides on soil microbial communities typically focus on pesticides in isolation [5, 22]. However, the long-term Nash’s Field grassland experiment demonstrated that interactions between pesticides and other stressors (e.g. pH) are essential for predicting shifts in soil bacterial community composition [9].

We investigate how combined fungicide exposure and warming temperatures affects diversity, composition and functioning of soil bacterial communities. Fungicides are the most widely sold class of pesticides in the EU [17], and their extensive use has led to frequent detections and residue accumulation in agricultural soils [3]. At the same time, global temperature projections indicate significant warming of soils (1°C to 6°C increase by 2100) [23]. Soil warming can significantly alter soil bacterial community diversity, composition, and functioning, favouring fast-growing heat tolerant taxa [24–26]. Moreover, warming can increase pesticide toxicity, as shown for fish, earthworms and mosquitos [27–29]. However, it remains unclear whether simultaneous warming and pesticide exposure may interact to shape non-target (i.e. bacterial) soil community composition, diversity and ecosystem functioning, as well as whether these changes matter for plants. To address this question, we conducted a fully factorial soil microcosm experiment where we manipulated exposure to fungicide (Fubol Gold, a mixture of mancozeb and metalaxyl-M) and/or warming temperatures over a period of six months. We assessed how fungicide and warming interacted to shape soil bacterial community composition, diversity, functioning (respiration and metabolic diversity), adaptation (fungicide resistance), plant growth promotion (Spring Barley, *Hordeum vulgare*, cv. Curtis) and herbivore population growth (*Sitobion avenae)*.

## 2. Methods

### 2.1 Community experimental evolution

a. *Extracting a soil community for experimental evolution:* A soil microbial community was extracted from John Innes No.2 compost (J. Arthur Bowers) by shaking 200g compost with 400 sterile glass beads in 200 mL M9 buffer in a sterile 1L Duran for 2min. After settling for 15 min, the supernatant was collected and cryopreserved in 20% (v/v) glycerol at −80 °*C*.
b. *Community experimental evolution:* Thirty-two sterilised soil microcosms were prepared by twice-autoclaving 30ml glass universal tubes containing 10g compost. Microcosms were left at room temperature for five days to allow reoxidation of sterilisation by-products [30], before re-adding 100µl extracted soil community (supplementary methods *SM1.1*). We chose to sterilise and re-add the resident community to maintain similarity between evolution conditions and later functional tests which require soil sterilisation. For fungicide treatments (n=16), we used “Fubol gold” (Syngenta; 64% w/w mancozeb + 3.88% w/w metalaxyl-M). Fubol Gold was dissolved in sterile water to a concentration of 1g/ml, before diluting to the relevant working concentration, which increased at 4-week intervals to simulate fungicide accumulation in soil (Weeks 1–4: 10 mg kg⁻¹, Weeks 5–8: 50 mg kg⁻¹, Weeks 9–12: 100 mg kg⁻¹, Weeks 13–16: 500 mg kg⁻¹). As a closed system, it was not our aim to mimic natural concentrations of fungicide found in agricultural soils, but to ensure constant strong directional selection for fungicide resistance throughout the experiment. Half of the microcosms (n=16) received 800µL fungicide solution, and half received 800 µL sterile water. Microcosms were vortexed for 30s and incubated statically (26 °C, 70% RH) for 16 weeks. Every 7 days, microcosms were washed with 10mL sterile M9, vortexed with 20 sterile 5-mm glass beads, before transferring 100 µL of soil wash to fresh sterile soil containing the appropriate fungicide concentration. Half of the microcosms were maintained at 26°C, while the remainder were subjected to a weekly temperature increase of 0.5 °C to a maximum of 34 °C, yielding four treatments (n = 8 each; 32 microcosms total). After 16 weeks, soil washes and soils were stored at −80 °C with or without 20% (w/v) glycerol, respectively.

### 2.2 Community 16S rRNA gene metabarcoding

To analyse taxa and diversity shifts in bacterial communities evolved under the different treatment conditions, we performed DNA extraction and metabarcoding sequencing. Soil from week 16 of the evolution experiment that had been frozen at -80 °C without glycerol was thawed, and DNA was extracted using a Qiagen DNeasy PowerSoil Pro Kit. DNA quantity and quality were measured using a nanodrop. Freezing samples at -80 °C without a cryoprotectant is commonly used for metabarcoding analysis and has a consistent or negligible effect on community composition [31–33]. Metabarcoding was performed by the Centre for Genomic Research (Liverpool, UK) using Illumina MiSeq (2 × 150 bp) targeting bacterial 16S rRNA genes (NimaGen 16S kit). Reads were demultiplexed, adapter-trimmed (Cutadapt v1.2.1), quality-checked (FastQC), and trimmed with Sickle (v1.200; Q ≥ 20, min length 15 bp). Sequences were denoised with DADA2 [34] within QIIME2, trimmed to 150 bp, and chimeras removed. Two samples containing only *Pseudomonas* were excluded. Diversity analyses used rarefied data (120,000 reads), retaining 83.1% of sequences and removing three samples (C5, W8, and WF8). Taxonomy was assigned using the SILVA 138 classifier, and phylogenetic analyses were conducted with MAFFT and FastTree [35].

### 2.3 Assessing community productivity

Respiration (CO_2_ production) was measured as a proxy for community productivity using MicroResp^TM^ [35]. Compost was sieved (∼ 1.2mm) and 0.35 g was added to all wells of a deep-well 96-well plate, covered with a gas-permeable metal lid, twice-autoclaved (126°C, 15min), and rested for five days. 50 µL sterile water was added to each well, vortexed for 30 seconds and left for 1h before community re-inoculation. Respiration assays and community metabolic profiling (see below) were run concurrently. The evolved and ancestral communities were prepared for these experimentsby inoculating 100µL thawed soil wash into 10g sterile soil and incubated under control conditions (no fungicide, 26°C) for 14 days, with transfer after seven days. After 14 days, 50µL soil wash was inoculated into two or three wells of a deep-well 96-well plate containing 0.35 g of sterilised soil (2 replicates per evolved community, 3 replicates for ancestral community), vortexed, and incubated at 26°C for seven days. Respiration was measured over 24h at 26°C using a colorimetric detection lid (Supplementary methods *SM1.2*), with OD_580_ recorded before and after incubation. ΔOD_580_ values were calculated, with higher values indicating greater respiration.

### 2.4 Community metabolic profiling

Metabolic profiles were assessed using Biolog EcoPlates. For each evolved community, 1mL soil wash was centrifuged (1 min, 500 g), and 800 µL supernatant was diluted 10× in sterile 0.85% NaCl. Aliquots (150 µL) were inoculated into each of 32 wells per plate (31 carbon sources, one water control) and incubated at 26°C for eight days. OD_580_ was measured on day 8 using a Tecan Nano plate reader. Carbon utilisation was quantified as ΔOD_580_ (final − blank), with higher values indicating greater metabolic activity. Evolved communities were assayed once; ancestral and uninoculated controls were run in triplicate.

### 2.5 Plant performance assays

We tested how each community influenced (i) barley (*Hordeum vulgare*, cv. Curtis) growth and herbivore population growth rates of a sap-feeding agricultural aphid pest (*Sitobion avenae*) [36]. Preserved soil washes were revived by inoculating 100µL into 10g sterilized soil and incubating 14 days, then added to plant pots with mesh lids (∼600mL pots; 32 evolved communities × 3 replicates = 96 pots). Surface-sterilized seeds were planted 1cm deep, watered (3mL) and grown at 26 °C, 60% RH, 16:8 light-dark. Aphids were pre-acclimated for 7 days, and 2 fourth-instar aphids were added on day 6. Plant height and adult aphid numbers were measured on days 16, and 20; aphids were counted on days 16 and 20. On day 20, roots and shoots were harvested, dried at 50 °C for 5 days, and weighed to determine dry biomass. The germination rate of our Spring barley seeds was ∼50%, which may be reflective of our experimental growth conditions which were optimised for the bacterial soil community rather than the plant (26 °C, rather than 15-20 °C) preferred by Spring barley [37]. As a result of this low germination rate, the following soil communities were excluded from the analysis due to complete failure of all 3 technical replicates to germinate: C7, F1, W1, and W6.

### 2.6 Quantifying fungicide minimum inhibitory concentrations (MIC’s) of evolved soil isolates

We chose 3 communities at random per treatment for fungicide resistance assays on six individual isolates from each community (n= 18 isolates per treatment; Supplementary methods *SM1.3*). Isolates were grown from frozen stocks overnight in LB broth, standardized to ∼1.5 × 10⁸ CFU/mL, and 10 µL added to each well of a 96-well plate containing Mueller-Hinton (MH) broth and x2 serial dilutions of Fubol Gold fungicide (increasing from 5 mg/mL). Plates were incubated statically at 28 °C for 18h. Three technical replicates were performed for each isolate. The MIC was taken as the minimum concentration (mg/mL) of fungicide which resulted in no detectible bacterial growth.

### 2.7 Statistical analysis

#### 16S rRNA gene metabarcoding

Weighted and unweighted UniFrac distance matrices were calculated from the feature table and phylogenetic tree. Differences in β-diversity were tested using PERMANOVA, with pairwise PERMANOVA for post hoc comparisons. α-diversity (Faith’s phylogenetic diversity and Pielou’s evenness) was analysed using linear models including warming, fungicide, and their interaction, followed by Holm-corrected t-tests to assess differences between treatments. Differentially abundant bacterial families were identified using ANCOM, with W values indicating the number of pairwise rejections of the null hypothesis.

#### Community productivity and metabolic capacity

Respiration and Biolog EcoPlate assays were established from the same inoculum. Six communities (C1, C3, C5, C8, WF3, WF8) failed to colonize fresh soil and showed no detectable respiration, metabolic activity, or growth on LB agar and were excluded from analyses. Two additional communities (W4, W7) were excluded due to absence of detectable bacterial communities in 16S data. Mean metabolic capacity was quantified as average well-colour development (AWCD), calculated from OD_580_ values across 31 substrates after subtraction of blank controls; values <0.06 were set to zero. Substrate richness was defined as the number of positive wells (OD_580_ > 0.06). Treatment effects on respiration (ΔOD_580_) and AWCD (day 8) were tested using linear models with warming, fungicide, and their interaction, with significance assessed via F-tests of nested models. Treatment specific differences were analysed using ANOVA with Tukey HSD. Overall metabolic profiles were compared using PERMANOVA (adonis2, vegan) based on Euclidean distances, while individual substrates were analysed using one-way ANOVA with Benjamini–Hochberg FDR correction and Holm-adjusted pairwise tests.

#### Plant and aphid performance

Five plants were excluded due to poor establishment (height <10 cm by day 16) or extreme outliers in root length. MANOVA tested treatment effects on shoot biomass, root biomass, and root length, followed by univariate ANOVAs. Aphid abundance (days 16 and 20) was analysed using negative binomial generalized linear models including treatments, their interaction, and plant height as a covariate; significance was assessed using likelihood ratio tests.

#### Fungicide resistance

Log_2_-transformed MIC values were analysed using linear mixed-effects models with fungicide, warming, and their interaction as fixed effects and community ID as a random effect; significance was assessed using likelihood ratio tests.

Statistical analyses were performed using R version 4.5.1 [38]

## 3. Results

### 3.1 Fungicide alters bacterial community composition and reduces diversity

Fungicide reduced bacterial α-diversity (Faith’s phylogenetic diversity; F_1,24_ = 309.05, p < 0.0001; Fig. 1a), whereas warming had no detectable effect (F_1,24_ = 0.44, p > 0.05). The reduction in diversity due to fungicide was not influenced by warming (F_1,23_ = 2.55, p > 0.05). Pairwise comparisons confirmed that fungicide, alone or with warming, decreased α-diversity relative to controls (p.adj < 0.001), while warming alone did not (p.adj > 0.05). Fungicide also reduced community evenness (Pielou’s index; F_1,24_ = 79.33, p < 0.0001; Fig. 1b). Warming alone had no significant effect on evenness (F_1,24_ = 3.35, p > 0.05), but partially mitigated the fungicide-induced decline in evenness (fungicide x warming, F_1,23_ = 69, p < 0.0001). Note that Pielou’s evenness does not account for phylogenetic relatedness among taxa, unlike Faith’s phylogenetic diversity. Our result provides strong support for fungicide-driven loss of bacterial diversity in soil communities.

**Figure 1.**
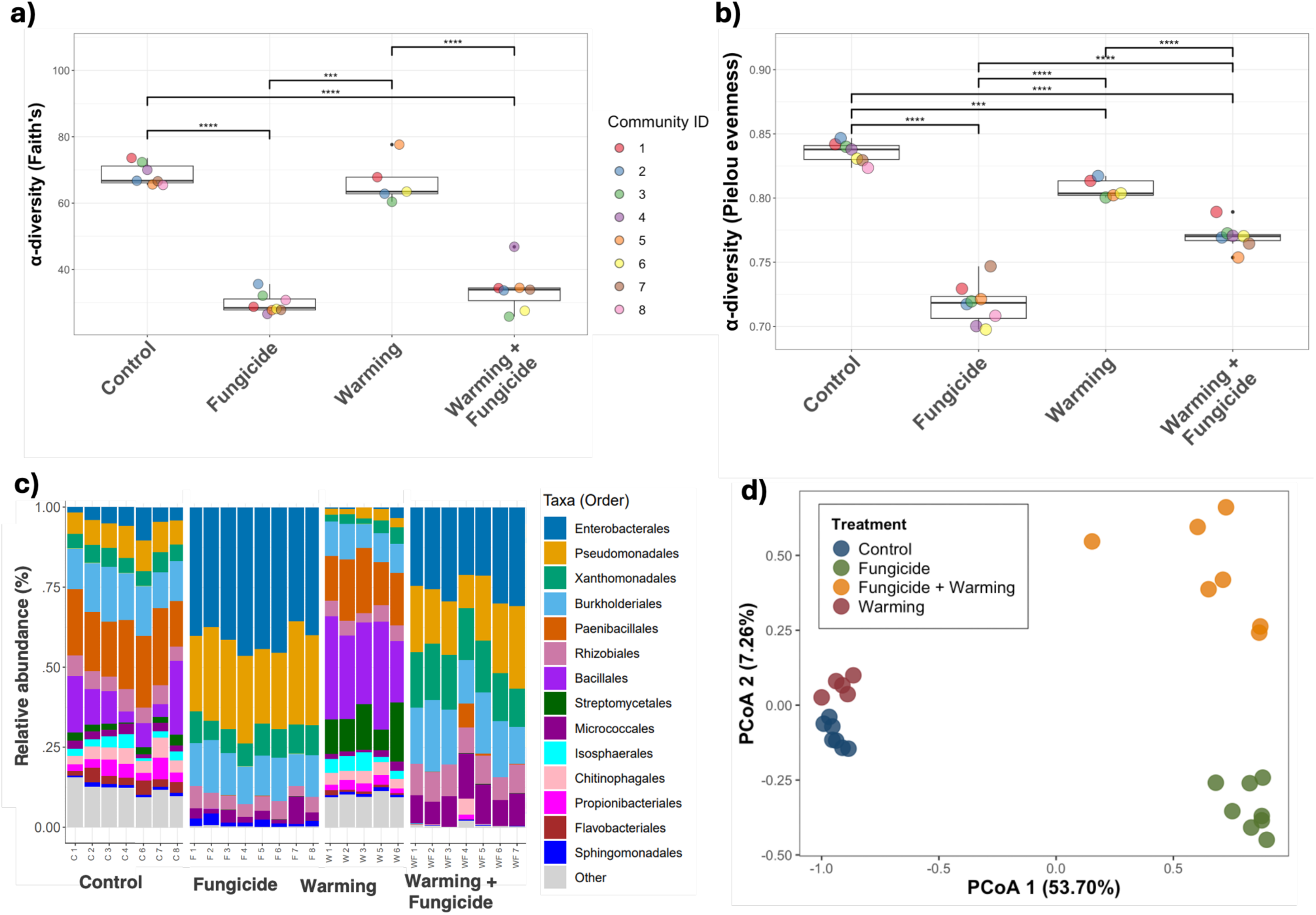
**(a–d).** Effects of warming and fungicide treatment on bacterial community diversity, assessed via 16S rRNA gene metabarcoding. **a.** Faith’s phylogenetic diversity was significantly reduced in fungicide-treated communities compared to controls, regardless of warming treatment. **b.** Pielou’s evenness was lower in all treated communities (warming, fungicide, or both) relative to controls, indicating that environmental stressors favoured dominance by a subset of bacterial taxa. Boxplots show the median (central line), interquartile range (box), and 1.5× interquartile range (whiskers). Individual points represent diversity values for each community and are color-coded by community ID. Lines connecting groups indicate significant differences based on pairwise t-tests with Holm correction. Significance levels are denoted as follows: p.adj < 0.05 (*), p < 0.01 (**), p < 0.001 (***), p < 0.0001 (****). **c)** Relative abundances (%) of each taxa (Order) in each evolved community. **d.** PCoA plot showing treatment-specific clustering, with the first axis (53.7% of variation) separating fungicide from non-fungicide communities.

We assessed how treatments shaped bacterial community composition using β-diversity (UniFrac distances). Treatments significantly affected composition in both unweighted (F = 13.51, p = 0.001, Fig 1c) and weighted analyses (F = 38.69, p = 0.001). All treatments differed from controls (pairwise PERMANOVA: warming F = 2.82, fungicide F = 25.5, multi-stressor F = 16.35; all p = 0.001), and both single-stressor treatments differed from the multi-stressor treatment (warming F = 12.42, fungicide F = 4.43; p <0.01). PCoA showed treatment-specific clustering, with axis 1 (53.7% variation) separating fungicide from non-fungicide communities (Fig. 1d), highlighting the key role fungicide had in shaping microbial community composition. Across 120,000 reads per sample assigned to ≤112 bacterial families, 40 (∼36%) differed significantly among treatments (ANCOM; Table S1). Fungicide was associated with increased relative abundances of *Pseudomonadaceae* and *Enterobacteriaceae*, while warming increased relative abundance of *Streptomycetaceae* and *Bacillaceae* compared to controls.

### 3.2 Effect of treatment on CO_2_ production

We measured CO₂ production as a proxy for community productivity for evolved and ancestral communities. Productivity was explained by a significant fungicide × warming interaction (F_1,44_= 9.01, p < 0.01; Fig. 2a), with dual-stressor communities showing lower CO₂ production than expected from single-stressor effects. Dual-stressor communities had reduced CO₂ relative to control, fungicide, and warming treatments (Tukey HSD, p < 0.001), while single stressors did not differ from controls (Tukey HSD, p > 0.05). The dual stressor treatment caused a synergistic reduction in CO_2_ production relative to stressors imposed independently (mean change in CO_2_ productivity caused by fungicide (+2.37%), warming (−10.9%) and multi-stressor treatment (−64.5%), compared to control). CO_2_ production under control, fungicide, and warming treatments exceeded that of the ancestor Tukey HSD, p < 0.0001, Fig 2a), whereas multi-stressor communities did not (Tukey HSD p > 0.05). Hence, interactions between temperature and fungicide have a distinct, non-additive effect on community respiration.

**Figure 2.**
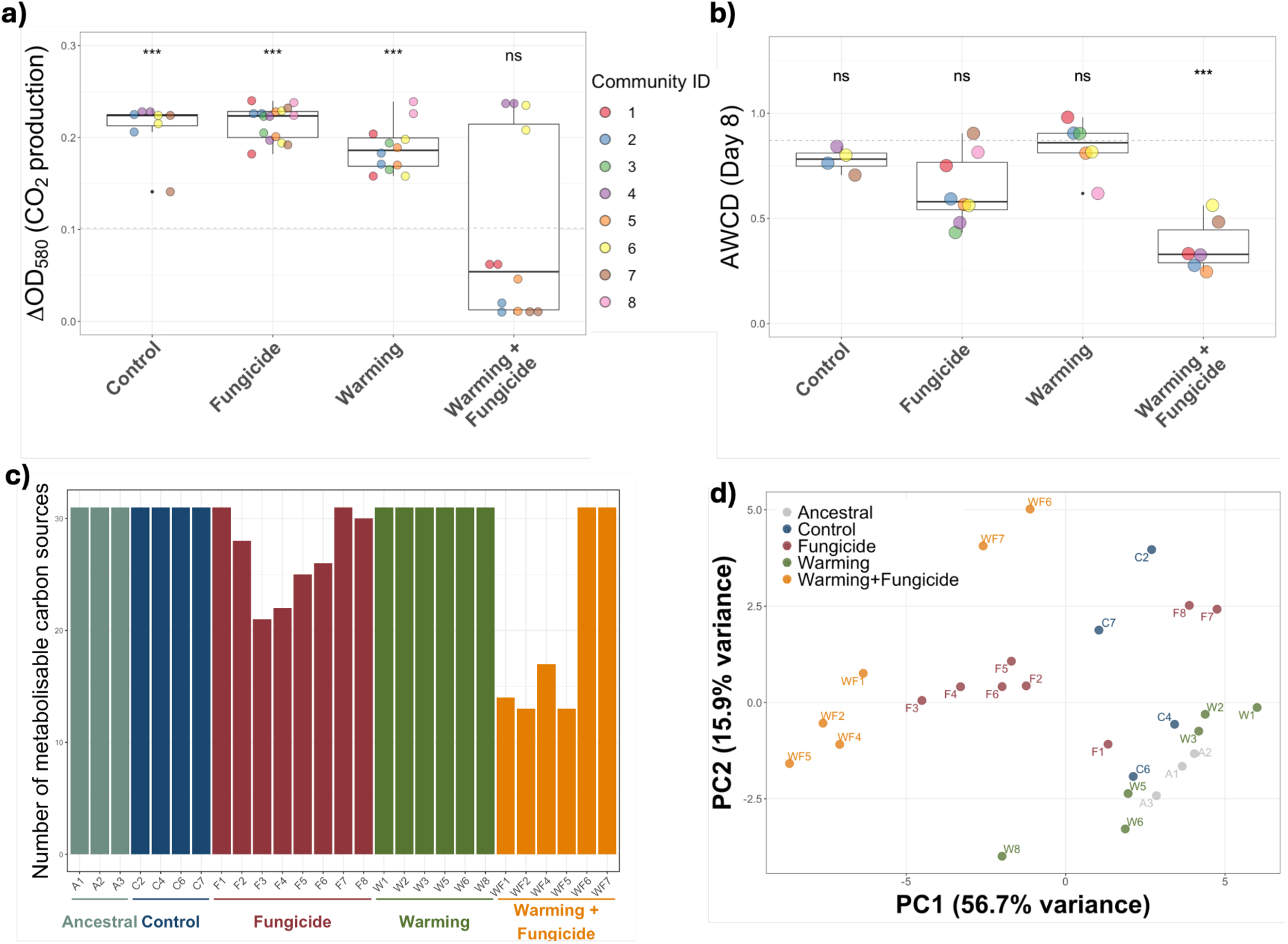
**(a–d).** Effects of warming and fungicide treatment on community functioning. **a.** Warming and fungicide treated communities have reduced CO_2_ production compared to single stressor effects alone (LM: fungicide x warming interaction). Dual-stressor communities had reduced CO_2_ production compared to control, fungicide, and warming communities, while no differences was observed between controls and single stressor treatments. Control, Fungicide and Warming treated communities have significantly greater CO_2_ production compared to ancestral communities, while dual stressor communities have, on average, similar CO_2_ production compared to ancestral communities. Asterisks indicate whether treatments differ significantly compared to ancestors (ns; no significant difference). **b.** Average well colour development (AWCD) is reduced in dual stressor communities, compared to the effects of fungicide and warming alone (LM fungicide x warming interaction). Dual-stressor communities had reduced CO_2_ production compared to control, fungicide and warming communities, while no differences was observed between controls and single stressor treatments. Boxplots show the median (central line), interquartile range (box), and 1.5× interquartile range (whiskers). Individual points represent values for each community and are color-coded by community ID. Significance levels are denoted as follows: p < 0.05 (*), p < 0.01 (**), p < 0.001 (***), p < 0.0001 (****). **c.** Number of metabolizable carbon sources (maximum 31) for evolved and ancestral communities**. d.** Treatment significantly affected community-level substrate utilization patterns. Dual stressor treatments clustered together on PC1 (orange points) with the exception of WF6 and WF7, which retained the ability to at least partly metabolise all 31 substrates.

### 3.3 Treatment-specific differences in community metabolic profiles

To investigate whether stress treatment alters community metabolic capacity, we analysed the ability of our evolved communities to metabolise 31 different carbon sources. We assessed mean global metabolic potential across the 31 substrates using average-well-colour-development at day 8 (AWCD). A higher AWCD value indicates a high mean value for substrate utilisation across all 31 sources. AWCD was explained by an interaction between fungicide and warming, where the dual-stressor communities had lower AWCD values than expected from single stressor effects (fungicide x warming, F_1,20_=8.41, p<0.01, Fig 2b). Dual-stressor communities had reduced CO_2_ production compared to control (TukeyHSD, p< 0.001), fungicide (TukeyHSD, p< 0.01) and warming (Tukey HSD, p < 0.0001) communities, while no differences was observed between controls and single stressor treatments (TukeyHSD, p > 0.05 in all cases). The combination of fungicide + warming caused a synergistic reduction in AWCD relative to stressors imposed independently, with a mean reduction in AWCD compared to the control of -17.99% in the fungicide treatment, +7.84% in the warming treatment, and -52.31% in the multi-stressor treatment. Neither the control, warming, nor fungicide alone, had significantly different AWCD values compared to ancestral communities (Tukey HSD, p >0.05), but dual-stressor communities had significantly lower AWCD relative to ancestor (Tukey HSD, p >0.001). Similarly, while control and warming communities retained some ability to metabolise all 31 substrates (i.e. *C_i_ - R* > 0.06 for all substrates), fungicide treated communities had a mean substrate utilisation number of 27 (without warming) and 15.5 (with warming) (Fig 2c). This pattern was primarily driven by loss of carbohydrate and carboxylic acid metabolism in the fungicide treated communities, especially when fungicide was applied alongside warming (Supplementary Table S2).

Treatment significantly influenced community-level substrate use (PERMANOVA, F_4,26_= 9.23, p = 0.001). Dual-stressor communities clustered on PC1 (56.7% variance), except WF6 and WF7, which retained partial metabolism of all 31 substrates (Fig. 2c). The greatest contributors to treatment-specific differences were D,L-a-Glycerol Phosphate (ANOVA; F_4_=15.14, p<0.001), N-Acetyl-D-Glucosamine OD (ANOVA; F_4_ = 11.69, p<0.001), D-Glucosaminic Acid OD (ANOVA; F_4_=49.55, P<0.001), Glycyl-L-Glutamic Acid (ANOVA; F_4_ = 9.05, p<0.001), a-Ketobutyric Acid (ANOVA; F_4_ = 9.5, p<0.001) and i-Erythritol (ANOVA; F_4_ = 11.38, p<0.001), with 20/31 substrates showing significant treatment-specific effects (Supplementary Figure S1, Supplementary Table S3). Differences were mainly due to reduced metabolic activity in dual-stressor communities versus warming, ancestral, and control treatments (Supplementary Figure S2). Overall, dual stressors caused a loss of metabolic activity, particularly for carbohydrates and carboxylic acids, highlighting treatment-specific impairment of community function.

### 3.4 Treatment-specific effects on plant growth promotion and aphid suppression

We tested whether soil communities evolved under fungicide and/or warming, differed in their ability to promote growth and aphid defence in barley. At day 20, plant performance (root length, shoot biomass, root biomass) was overall shaped by a fungicide x warming interaction (MANOVA fungicide x warming interaction, Pillai’s trace = 0.44, F₃, ₂₁ = 5.49, p < 0.01). Univariate follow-up analyses showed that this interaction was driven primarily by effects on root length (F₁,₂₃ = 4.64, p <0.05), with a weaker, marginal effect on shoot biomass (F₁, ₂₃ = 4.15, p = 0.05), while root biomass was unaffected (p = 0.91). In both root length and shoot biomass, fungicide application under elevated temperature reduced plant performance relative to expectations from main effects alone (Fig. 3).

**Figure 3.**
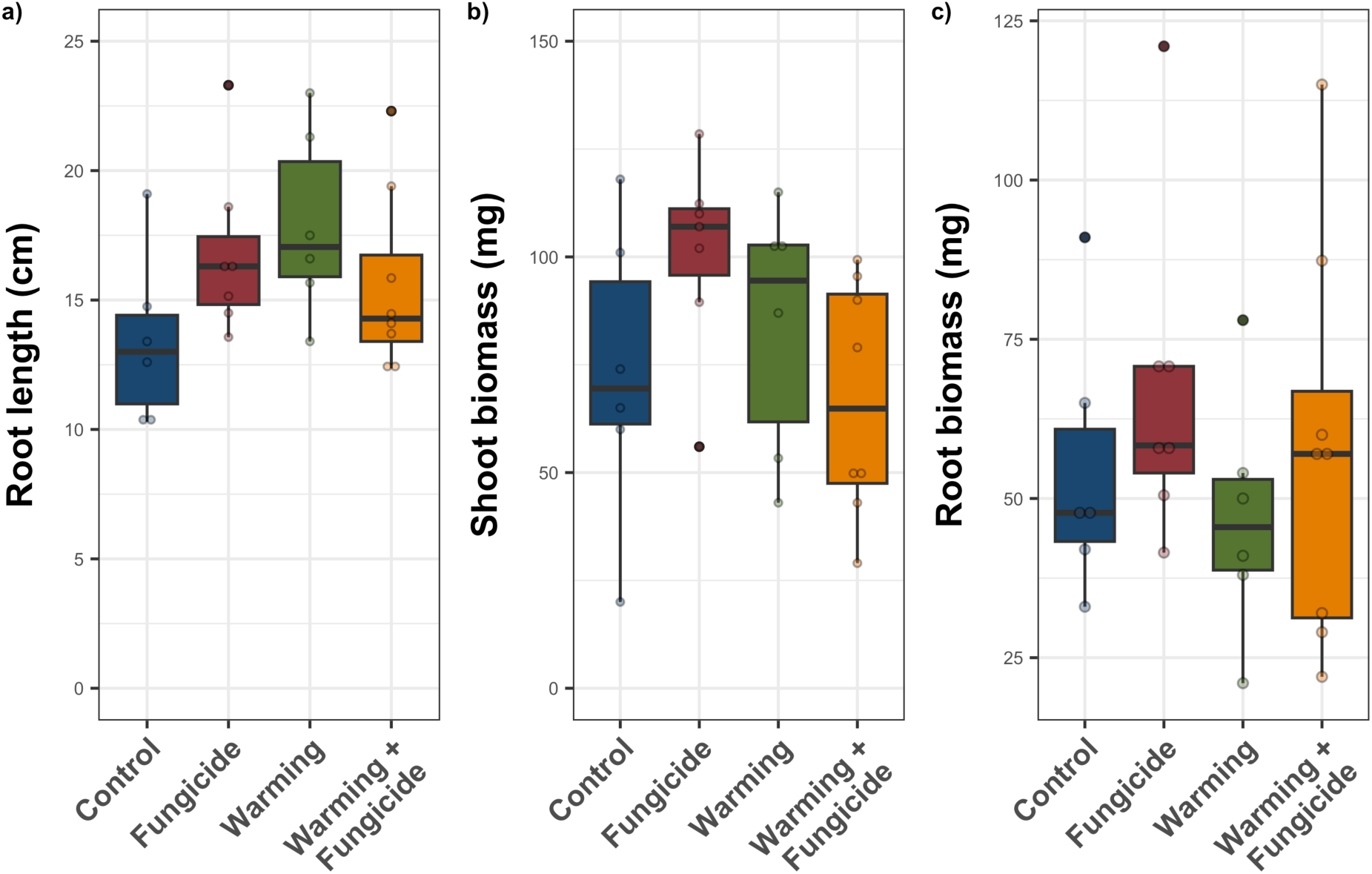
To assess plant growth promotion, barley seeds were grown in sterile compost inoculated with each evolved microbial community. On day 20, **(a)** root length was measured, and shoots and roots were harvested, dried, and weighed to determine **(b)** dry shoot biomass and **(c)** dry root biomass. Plant performance (root length, shoot biomass, root biomass) was overall shaped by a fungicide x warming interaction (MANOVA Pillai’s trace = 0.44, p < 0.01), driven primarily by effects on root length (p <0.05), with a weaker, marginal effect on shoot biomass (p = 0.05), while root biomass was unaffected (p = 0.91). In both root length and shoot biomass, fungicide application under elevated temperature reduced plant performance relative to expectations from main effects alone. Each datapoint represents the mean of 1–3 replicates per community. Boxplots show the median (central line), interquartile range (box), and 1.5× interquartile range (whiskers).

At day 16, fungicide treatment was associated with a marginal, but non-significant reduction in aphid abundance (χ² _1,23_= 3.72, *p*= 0.067), with no evidence that this effect depended on warming (fungicide × warming interaction: χ²_1,22_ = 1.04, *p* > 0.05) (Supplementary Figure S3). By day 20, the effect of fungicide was no longer detected (χ²_1,23_ = 1.14, *p* >0.05), and there was again no evidence of an interaction between fungicide and warming (χ²_1,23_= 0.54, *p* > 0.05). Warming alone had no detectable effect on aphid abundance at either time point (day 16: χ² _1,23_= 0.09, *p* > 0.05; day 20: χ²_1,23_ = 0.78, *p* > 0.05).

### 3.5 Quantifying fungicide resistance among soil isolates

We retrieved isolates from three randomly-selected communities from each treatment by spreading samples on a variety of agar types (Supplementary methods SM1.3) and selecting colonies after 7 days incubation at 28°C. Colonies were purified by re-isolation and screened for their ability to grow in media spiked with increasing concentrations of Fubol Gold, i.e. a fungicide minimal inhibitory concentration (MIC assay). We find that fungicide resistance is best explained by an interaction between fungicide and warming, so that soil isolates were more resistant to fungicide only when fungicide was applied as a single stressor (i.e. in the absence of warming) (LMER; fungicide x warming interaction, χ^2^_1,5_ = 5.04, p <0.05, Fig. 4). For one of our randomly selected communities (W7), our 16s sequencing could not detect a community, so we re-ran our analyses with W7 omitted and found the same interaction (LMER; fungicide x warming interaction, W7 omitted, χ^2^_1,5_ = 4.2, p <0.05). The mode MIC for isolates from fungicide-evolved communities was 16-fold higher relative to the control (0.0097 mg/mL versus 0.156 mg/mL), suggesting fungicide drives selection for fungicide resistance. Warming also increased mode MIC 4-fold relative to control isolates. However, despite both stressors independently selecting for resistance, the mode MIC of isolates from dual stressor communities did not differ from control isolates (0.0097 mg/mL). Hence, fungicide can select for soil isolates that are significantly more resistant to the fungicide, but selection for resistance is constrained in the dual-stressor environment.

**Figure 4:**
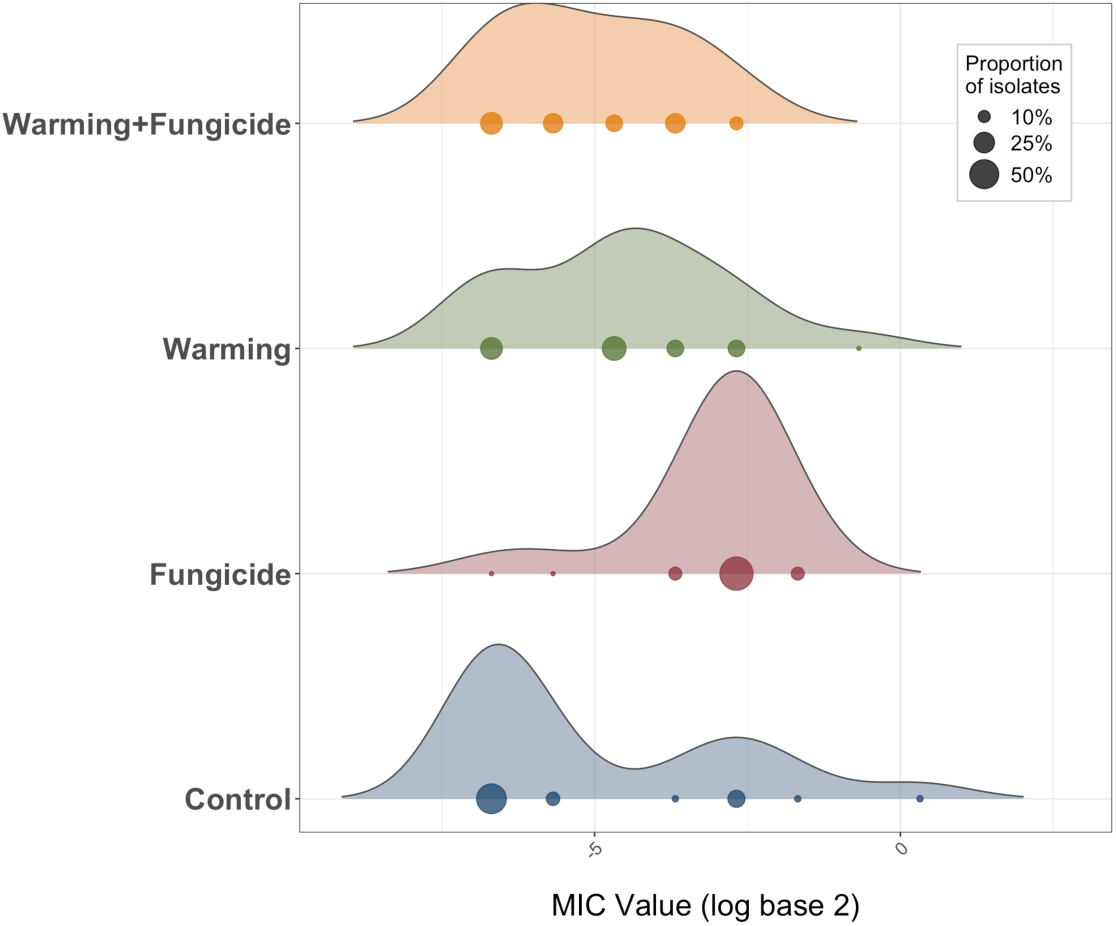
72 soil isolates (18 isolates per treatment) were screened for their ability to grow in media spiked with increasing concentrations of Fubol Gold, starting from 5 mg/mL. The MIC was taken as the minimum concentration (mg/mL) of fungicide which resulted in no detectible bacterial growth; X-axis shows mode log_2_ MIC for each isolate. Circles are scaled by the proportion of isolates displaying a particular MIC. Fungicide resistance was best explained by an interaction between fungicide and warming, so that soil isolates were more resistant to fungicide only when fungicide was applied as a single stressor.

## 4. Discussion

We explored how two environmental stressors (fungicide and warming temperatures) interact to shape soil bacterial community diversity, composition, adaptation, and functioning. Fungicide was a key driver of bacterial community composition, shifting communities toward Gram-negative taxa such as Burkholderiales, Xanthomonadales, Pseudomonadales, and Enterobacterales, while reducing overall diversity. Stressors imposed individually versus simultaneously had distinct impacts on community function (respiration rates and metabolic capacity), with dual-stressor communities exhibiting significantly reduced functioning compared to predictions from individual effects. Dual-stressor communities lost the ability to metabolise a range of carbon sources, particularly carbohydrates and carboxylic acids, to a much greater extent than single-stressor communities. Moreover, isolates from dual-stressor communities did not display the increased levels of fungicide resistance observed in fungicide-only treatments, suggesting that warming impeded the emergence of resistant isolates. Finally, plant experiments revealed that while soils treated with warming or fungicide independently improved plant growth promotion, this effect was lost when stressors were applied simultaneously. Together, these results demonstrate that combined environmental stressors can reduce diversity, constrain adaptation, alter community composition, and impair functional capacity beyond the sum of individual effects.

In our study, fungicide accumulation under warming temperatures synergistically reduced community productivity (CO₂ production) and metabolic diversity compared to stressors imposed in isolation. Few studies have directly tested the combined impacts of fungicide and warming on soil communities. Drocco et al. [22] exposed soil microcosms to environmental disturbance followed by pesticide application and reported strong effects of disturbance on community diversity and structure, whereas pesticide effects were comparatively weak. In contrast, we observed strong interactive effects, likely because stressors were imposed simultaneously rather than sequentially as in [22]. Our finding that dual-stressor communities exhibited greatly reduced CO₂ production and metabolic capacity may reflect each stressor selecting for different taxa, such that their combination creates sub-optimal conditions for surviving communities. Consistent with this, isolates from dual-stressor communities showed no evidence of fungicide adaptation, unlike the 16-fold increase observed in fungicide-only evolved communities. Constraints on adaptation in multistressor environments are common in evolution experiments [39–41], although resistance in our study may also have arisen via species sorting toward resistant taxa, with both processes likely contributing [42].

In our study, fungicide accumulation under warming temperatures synergistically reduced community productivity (CO_2_ production) and metabolic diversity compared to stressors imposed in isolation. Few studies directly test the impact of fungicide and/or warming temperatures on soil communities. Drocco et al [22] exposed soil microcosms to an environmental disturbance (heat, high humidity, or no disturbance), and after 3 days of recovery, exposed microcosms to pesticides. The study reported a clear effect of environmental disturbance on community diversity and structure, but pesticide effects were sparse and much weaker in comparison, in contrast to the interactive effects we observed in our experiment. However, Drocco et al [22] staggered stressors, while we imposed stressors simultaneously. Our finding that dual-stressor communities exhibited greatly reduced CO₂ production and metabolic capacity may reflect each stressor selecting for different taxa, such that their combination creates sub-optimal conditions for surviving communities. Consistent with this, isolates from dual-stressor communities showed no evidence of fungicide adaptation, unlike the 16-fold increase observed in fungicide-only evolved communities. Constraints on adaptation in multistressor environments are common in evolution experiments [39–41], although resistance in our study may also have arisen via species sorting toward resistant taxa, with both processes likely contributing [42].

We found that bacterial communities had reduced α-diversity (i.e., richness and evenness) under long-term exposure to fungicides. In contrast, a meta-analysis of 73 studies comparing soil microbial communities between fungicide-free and fungicide-treated soils, found no detectible effect of fungicides on bacterial community diversity [3]. However, it is unclear how many of the included studies involved long-term fungicide exposure (i.e. >7 days), so we cannot directly contextualise our results within the scope of this meta-analysis. Fungicide exposure increased the relative abundance of Gram-negative taxa, including Burkholderiales, Xanthomonadales, Pseudomonadales, and Enterobacterales, while decreasing Gram-positive groups such as Streptomycetales, Paenibacillales, and Bacillales. Increases in Pseudomonadota following fungicide application have been reported previously [43], including metalaxyl–mancozeb treated soils [44], and Enterobacterales may possess pesticide-degrading capabilities [45, 46]. Notably, Gram-negative taxa are more prone to multi-drug resistance [47], raising the question of whether fungicide-treated soils represent an increased risk of harbouring resistant bacteria.

The composition of the barley root microbiome, which is recruited by plants from bulk soil, is known to influence both plant growth and aphid suppression [48]. We found that plants grown in soils with a history of fungicide or warming typically exhibited increased root length and shoot biomass, despite slightly reduced respiration rates and metabolic capacity in fungicide-treated soils. This beneficial effect may be due to fungicide targeting plant fungal pathogens (although we did not characterise the fungal community in this study). In contrast, when fungicide and warming were imposed simultaneously, this growth promotion effect of fungicide was lost. In our study, plants were grown under common garden conditions (in absence of fungicide or warming), suggesting these stress “legacy effects” can influence aboveground processes. Such legacy effects of stress have been reported previously for soil microbial communities [4], where ecological memory of soil can affect ecosystem resilience.

Although the individual effects of fungicide and warming on soil communities have been examined across a range of experimental systems, soil types, warming regimes, and pesticide classes [22, 49–51], evidence increasingly suggests that the number of stressors has a greater influence on soil ecosystem functioning than the identity of individual stressors [4, 8]. Experimental constraints associated with multi-stressor designs remain a major challenge. Hierarchical approaches that prioritise stressors with the largest predicted impacts, combined with random partition designs (e.g. [52]) may help disentangle interactive effects. Integrating laboratory microcosms with field-based manipulations across diverse soil types will further improve ecological realism and help guide management strategies to maintain soil health under global change.

## Funding Information

The authors would like to acknowledge the financial support of NERC (MK: NE/S00713X/1), BBSRC Discovery (SOB: BB/T009446/1) and IRC New Foundations and Community Foundation Ireland (SOB: NF/2022/39250777).

## Notes

### Competing Interest Statement

The authors have declared no competing interest.

