## Supplementary for "Fungicide and warming interact to reduce soil ecosystem functioning"

**Supplementary Figures and Tables**

**Supplementary Table S1:** The relative abundances (green bars) of taxa in the table below (family) differed significantly between treatments. W = ANCOM statistic indicating the number of times the null hypothesis of no treatment-specific differences was rejected. Fungicide treated communities are increasingly dominated by Gram-negative bacteria, such as *Pseudomonadaceae* and *Enterbacteraceae.*

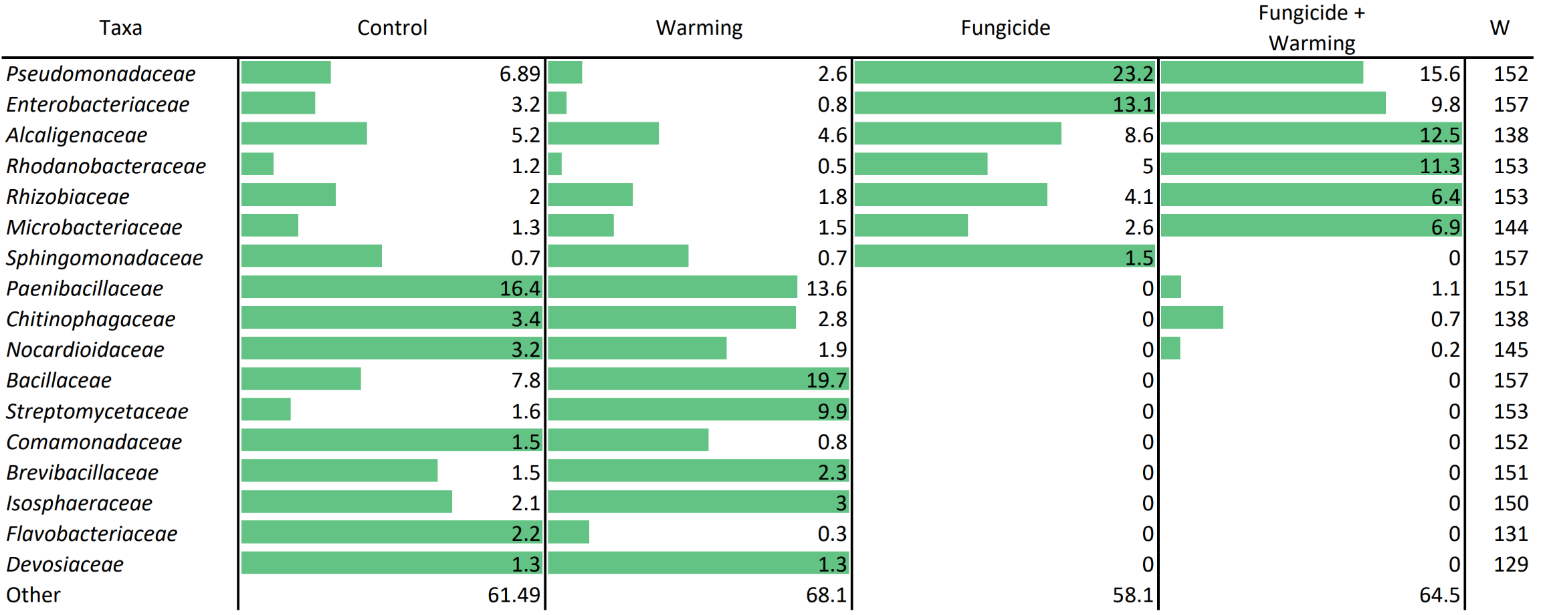

**Supplementary Table S2.** To assess whether stress treatments affect community metabolic capacity, we evaluated the ability of evolved microbial communities to metabolize 31 different carbon sources over an 8-day incubation period. The table presents data from day 8, showing the proportion of communities within each treatment group that retained the ability to metabolize each carbon source (defined as Ci - R > 0.06). All ancestral, control, and warming-evolved communities were able to metabolize all substrates. In contrast, fungicide-evolved communities—and more notably, those evolved under combined fungicide and warming treatments—exhibited widespread reductions in metabolic capacity, particularly for carbohydrates and carboxylic acids.

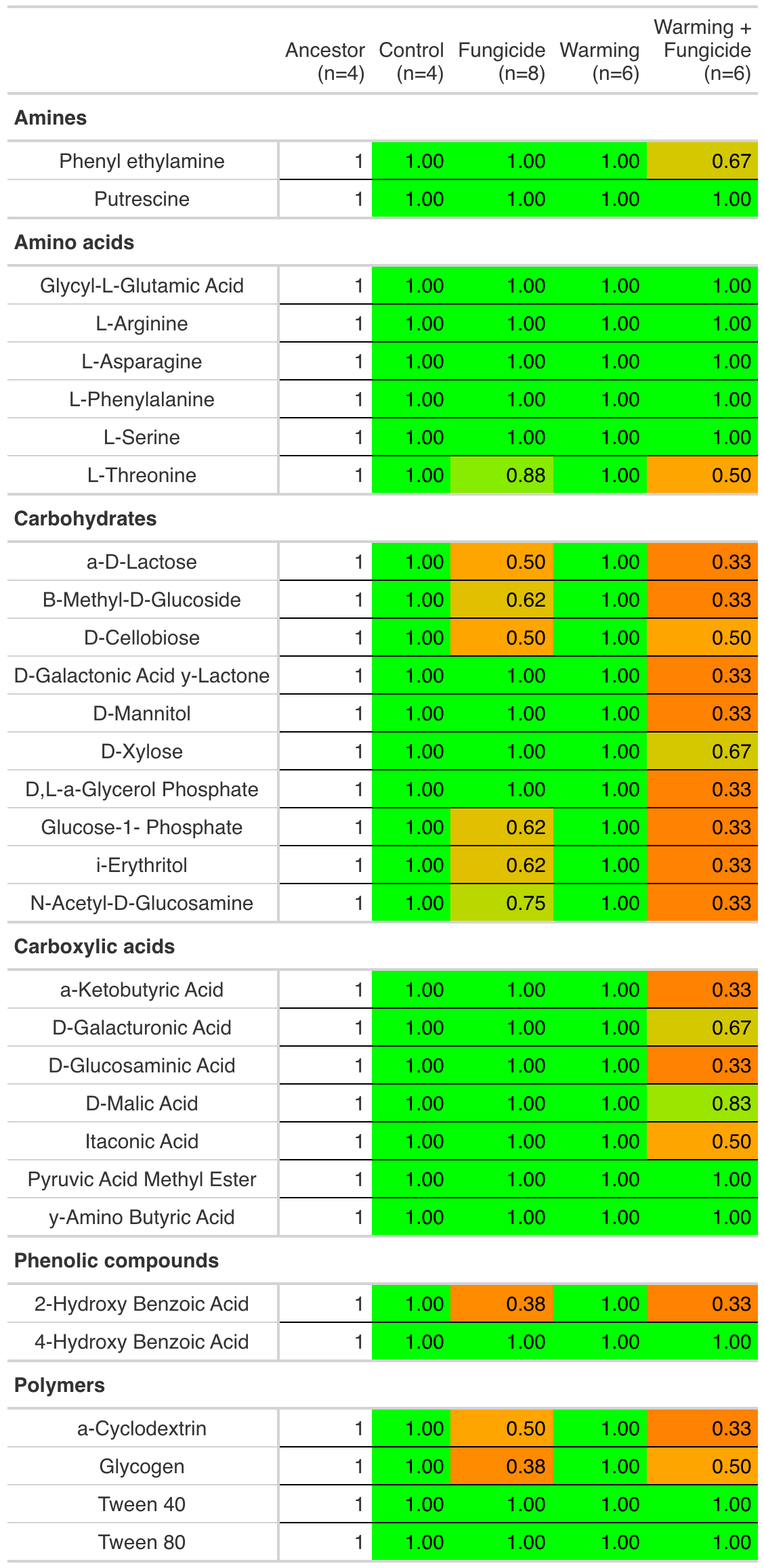

**Supplementary Table S3:** For each carbon source we quantified loadings on PC1 and tested each carbon source for treatment-specific differences using ANOVA with Benjamini-Hochberg correction and Tukey post-hoc tests. Carbon sources are listed in the order of their greatest contributors to treatment-specific differences (highest- lowest). Significant treatment-specific differences were found for 20 out of 31 carbon sources.

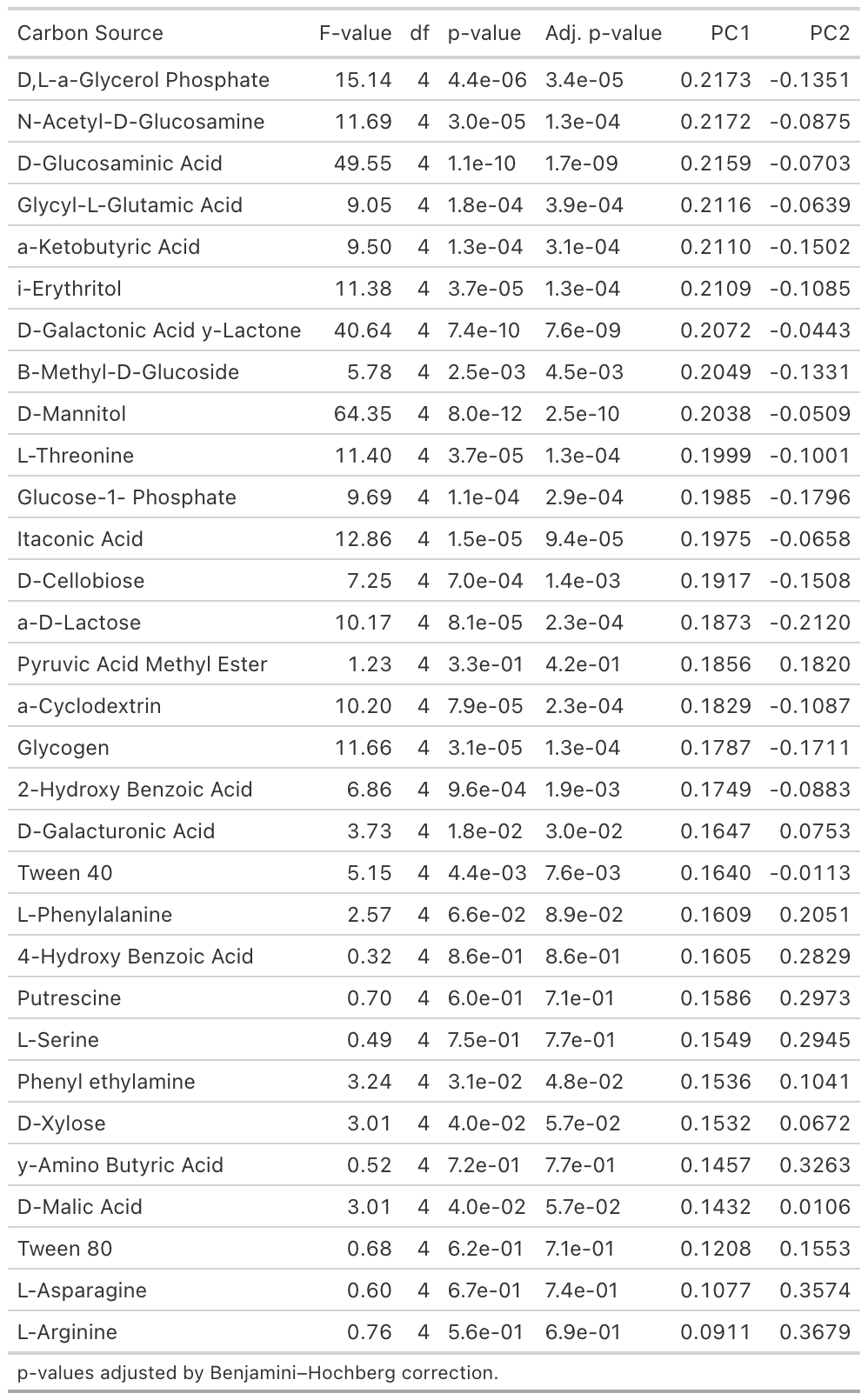

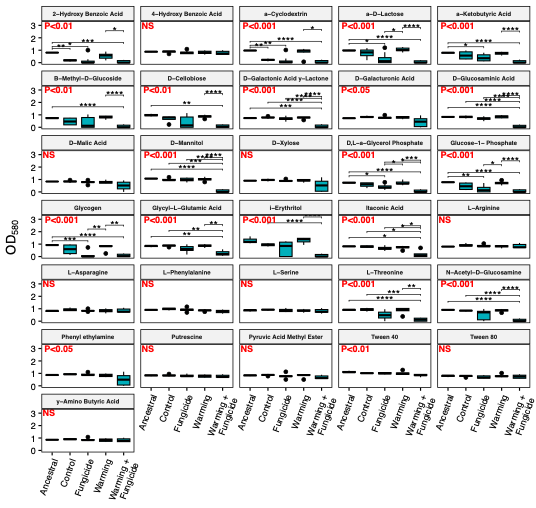

**Supplementary Figure S1:** For each Biolog carbon source ,we quantified treatment-specific differences in metabolic activity (OD580) using ANOVA with Benjamini-Hochberg correction and Tukey post-hoc tests. We see widespread loss of metabolic capacity in dual-stressor treatments (Warming + Fungicide) for 20 out of 31 carbon sources. ANOVA results (adjusted p-values) are shown in red. Significant post-hoc comparisons are shown as black connecting lines. Significance levels are denoted as follows: p < 0.05 (*), p < 0.01 (**), p < 0.001 (***), p < 0.0001 (****). Boxplots show the median (central line), interquartile range (box), and 1.5× interquartile range (whiskers).

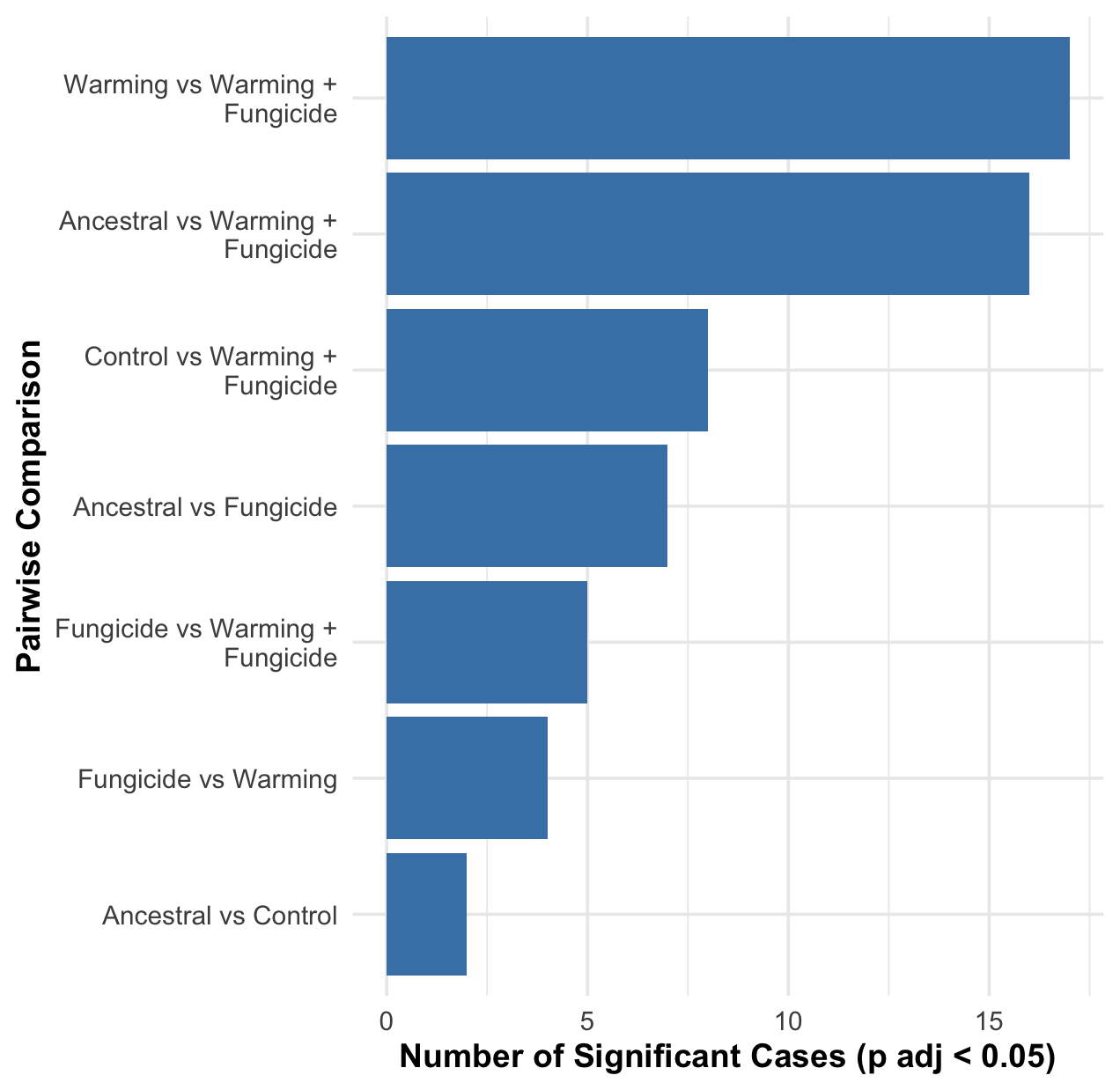

**Supplementary Figure S2**. Treatment-specific differences in metabolic activity were found for 20 out of 31 Biolog carbon sources. To identify the dominant treatment-level comparison driving these effects, we show the number of significant cases for each comparison across all 31 carbon sources. The dominant comparisons driving treatment-specific effects were reduced activity in warming + fungicide treatments versus warming (17/31 carbon sources), ancestral (16/31 carbon sources), and control (8/31 carbon sources) communities.

**b)**

**a)**

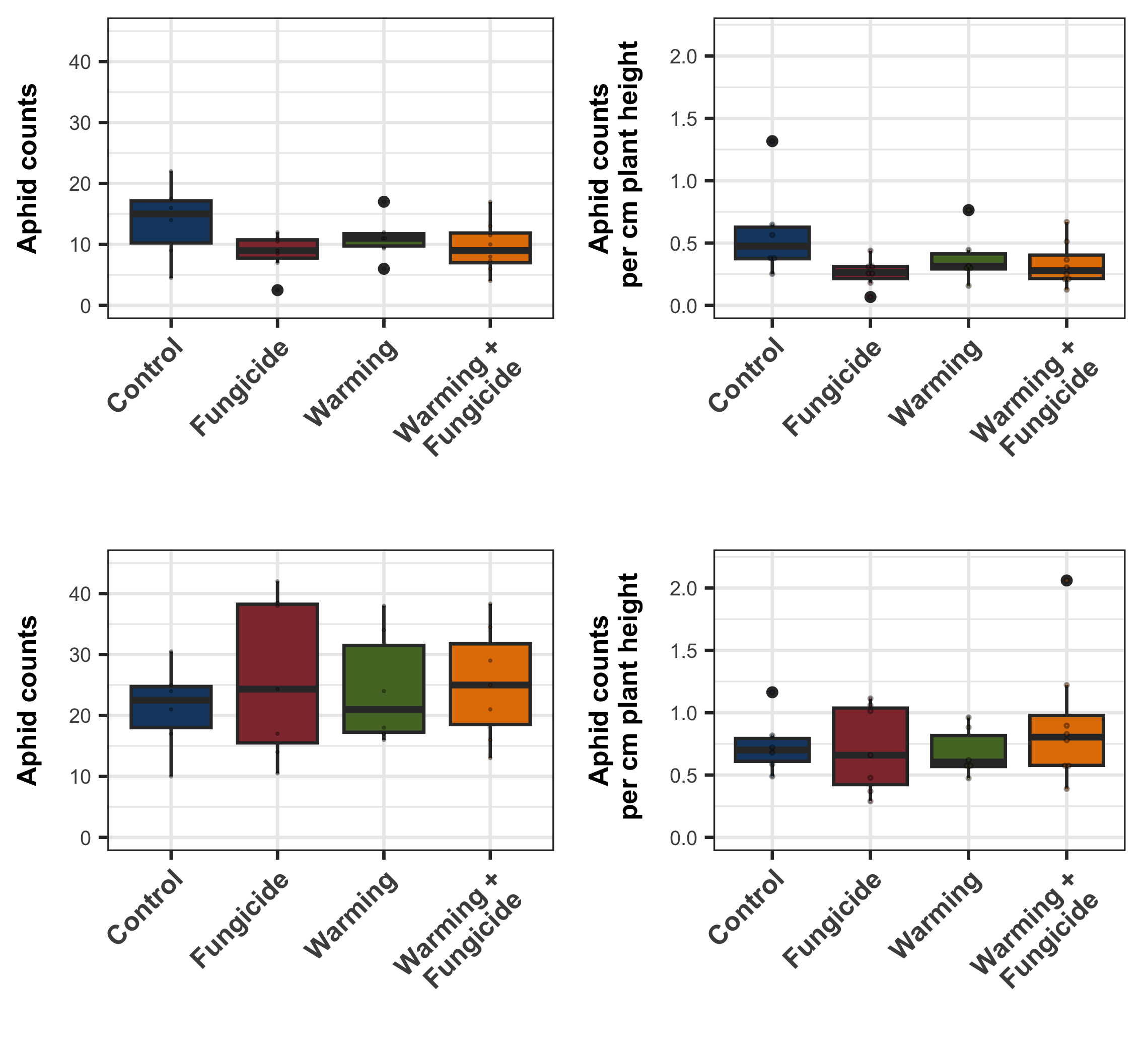

**DAY 16**

**DAY 16**

**DAY 20**

**DAY 20**

**d)**

**c)**

**Supplementary Figure S3:** Aphid abundance on barley plants grown in sterile compost inoculated with evolved microbial communities. Barley seeds were inoculated at planting, and aphids were introduced on day 6. Adult aphids were counted on days 16 (a, b) and 20 (c, d). At day 16, there was marginal but non-significant evidence that fungicide treatment reduced aphid abundance (χ²_1,23_ = 3.72, *p* = 0.067), but this effect was not detected by day 20. Temperature treatment had no detectable effect on aphid abundance at either time point, either alone or in interaction with fungicide. Each datapoint represents rounded mean aphid number per soil community (1–3 replicates per community). Boxplots show the median (central line), interquartile range (box), and 1.5× interquartile range (whiskers).
