## Supplementary methods for "Fungicide and warming interact to reduce soil ecosystem functioning"

*SM1.1 Community experimental evolution*

To our community microcosms, we note that we also added 100 µl (10^5^ CFU/ml) of *Pseudomonas fluorescens* tagged with a gentamicin-resistant cassette (SBW25::GentR) to each microcosm at a 1:1 ratio with the community, as we wanted to simultaneously investigate *de novo* adaptation of a focal species *P. fluorescens,* to fungicide and warming. However, this strain was quickly outcompeted by the community and could not be retrieved from the community by the end of the 16 week experiment.

*SM1.2 Preparation of respiration detection lid*

The respiration detection lid was prepared as follows. Indicator agar was produced by first preparing an indicator solution containing 16.77 g KCl (Sigma-Aldrich), 18.75 mg Cresol Red (Acros), and 0.315 g NaHCO_3_ (Sigma) in 1 l of distilled water heated to 50 °C in a water bath. A solution of 3% (w/v) Nobel agar (Sigma-Aldrich) was prepared in distilled water and heated in a microwave until completely dissolved. Agar was left in a water bath to cool to 50 °C, and then indicator solution was added to the agar in a 2:1 ratio (Indicator:agar) and gently stirred until homogenised. 150 µl of agar was added to each well of a 96-well plate (Thermo Scientific) and left to solidify. To standardise the OD of agar in each well, plates were sealed in an anaerobic jar containing soda lime (Thermo Scientific) and a beaker of water. After 48 hours, the plates were sealed with parafilm and returned to the anaerobic jar. The OD (580 nm) of each well was measured at the start and end of the experiment using a Tecan Nano Plate reader.

*SM1.3 Colony isolation for fungicide MIC assay*

We chose 3 communities at random per treatment for fungicide resistance assays on individual isolates: Control (C4, C5, C7), Fungicide (F1, F2, F5), Warming (W1, W2, W7) and Warming+Fungicide (WF1, WF4, WF5). Frozen community stocks were re-established in 10g fresh sterile soil for 7 days. A soil wash was extracted in PBS, diluted, and plated on Lysogeny broth agar (LB; Fisher Scientific), Reasoner's2A agar (R2A; Fisher Scientific), and Tryptone Soy Agar (TSA; Fisher Scientific) plates. Plates were incubated at 28°C for 7 days. A variety of agar types were used to isolate a variety of bacteria from each of the evolved communities, rather than selecting for bacteria that are fast growing and only grow on one type of agar (Supplementary Data 1).
